# Static and dynamic analysis of in-vivo imaging for early-detection of lymphedema via near-infrared fluorescence indocyanine green lymphangiography

**DOI:** 10.1101/2023.02.23.529802

**Authors:** Hwayeong Cheon, Sang-Hun Lee, Sang Ah Kim, Bumchul Kim, Hyunsuk Peter Suh, Jae Yong Jeon

## Abstract

**Background:** Near-infrared fluorescence indocyanine green (NIRF-ICG) lymphangiography, a primary modality for detecting lymphedema, which is a disease due to lymphatic obstruction, enables real-time observations of lymphatic flow and reveals not only the spatial distribution of drainage (static analysis), but also information on the lymphatic contraction (dynamic analysis).

**Methods:** We have produced lymphatic obstruction models in upper limbs through the dissection of proximal lymph nodes (LNs) and radiation (dissection limbs). After the model formation during 1 week, the static and dynamic analysis using NIRF-ICG lymphangiography were performed for six weeks. The drainage pattern and leakage of lymph fluid were observed and time-domain signals of lymphatic contraction were measured in the distal lymph vessels. The obtained signals were converted to the frequency-domain spectrums using the signal processing.

**Results:** The results of both static and dynamic analyses proved to be effective in accurately identify the extent of lymphatic disruption in the dissection limbs. The static analysis showed abnormal drainage patterns and an increased leakage of lymph fluid to the periphery of the vessels compared to the control limbs. Meanwhile, the waveforms were changed and the frequency of the contractile signals was increased by 58% in the dynamic analysis. Specifically, our findings revealed that regular lymphatic contractions, observed at a frequency range of 0.08 ∼ 0.13 Hz in the control limbs, were absent in the dissection limbs. The contractile regularity was not fully restored until the end of the follow-up, indicating a persistent lymphatic disruption in the dissected limbs.

**Conclusion:** The dynamic analysis was consistent with the static analysis, and it could detect the abnormalities of lymphatic circulation by observing the characteristics of signals without the need for a control group. As NIRF-ICG lymphangiography is currently used in clinical practice, our findings may be useful for the early detection of the lymphatic circulation problem.

## INTRODUCTION

Lymphedema, or lymphoedema, is a chronic disease caused by the disruption of the lymphatic circulation and accumulation of extracellular fluid, containing various substances, in the interstitial tissue near to the lymphatic vessels or lymph vessels [^1–5^]. Especially, cancer treatments involving LN dissection and radiotherapy substantially lead to an acquired defect and a physical block of lymphatic system and they may induce secondary lymphedema (acquired lymphedema). If this condition is not properly managed at an early stage, irreversible tissue transformation can occur such as the swelling of the extremities, tissue fibrosis, repeated cellulitis, and excessive adipose deposition [^6–8^]. The principal symptom of lymphedema is swelling, as indicated by the inclusion of “edema” in its title, and currently, the physical examinations such as measuring the circumference of extremities, observing the skin change appearance, or Stemmer’s sign are mainly performed in clinical settings. However, because the development of lymphedema begins before there are externally identifiable limb volume changes caused by the disruption of lymphatic transportation, this disorder can be diagnosed at an early stage only when the change of circulation is detected before these external changes occur. It is typically the case across many diseases that the earliest possible diagnosis and treatment are important for improving the prognosis. In the case of lymphedema, better outcomes are obtained when a diagnosis is made, and treatment commences, before physical change appears [^9–13^].

More effective detection techniques that can accurately evaluate lymphatic circulation are required for the earlier diagnosis and treatment of lymphedema. Lymphoscintigraphy and near-infrared fluorescence indocyanine green (NIRF-ICG) lymphangiography are the most well-known methods of visualizing and evaluating lymphatic flow and involve the injection of nuclear and fluorescent agents, respectively [^14–17^]. Among its advantages, NIRF-ICG lymphangiography or lymphography enables the real-time observation of indocyanine green (ICG) fluorescent molecules that have been injected into the interstitial tissue and been collected by lymphatic capillaries before transitioning along the small lymphatic vessels. Because ICG molecules mainly absorb infrared light at a 789-nm peak and emit fluorescence of an 814-nm peak, it is possible to observe lymphatic vessels and drainage on the skin without incision by using near-infrared light that penetrates the skin [^18,19^]. Although NIRF-ICG lymphangiography is usually used to observe superficial lymphatics due to penetration depth limitations and scattering by the dermis layers, it offers not only observations of the spatial distribution of lymphatic vessels, which are represented by the drainage pattern in static analysis, but can also dynamically analyze lymphatic pumping or lymphatic contraction in the form of time-domain signals [^20^]. In addition, because the physical quantities (frequency, waveform shapes, etc.) of lymphatic contractions in the time domain provide important information on the function of induvial lymphatic vessels, it may also be an effective technique for the preemptive evaluation of abnormalities in the lymphatic circulation before they start to affect the surrounding tissues. In addition, the frequency components of the periodic waveforms in the time domain can be analyzed by transform signal processing. In the frequency-domain spectrum, the characteristics of lymphatic contraction, which are described using only a single waveform in the time domain, can be further characterized by the sum of multiple independent frequency components.

In our present study using lymphedema animal models, the in vivo signal characteristics of lymphatic contraction in lymphedema were observed using the dynamic analysis properties of NIRF-ICG lymphangiography and the results were compared with those obtained via the static analysis that is usually used in clinical practice [^21–23^]. We induced the lymphatic disruption in the proximal area of in a limb of the animal models (dissection limbs), and the development of lymphedema in the limbs was evaluated using the volume differences with control limbs. In the NIRF-ICG lymphangiography experiments, static analysis for lymphatic drainage patterns and lymph fluid leakage was conducted in the dissection limbs in comparison to the control limbs. Dynamic analysis of the distal lymph vessels was also performed to observe any changes in lymphatic contraction. After converting the time-domain signals to the frequency-domain spectrum via two kinds of spectral analysis methods, fast Fourier transform (FFT) and continuous wavelet transform (CWT) [^24–26^], the frequency components of lymphatic contraction were analyzed mathematically.

## METHODS

### Experimental preparation

All animal procedures were approved by the Institutional Animal Care and Use Committee (IACUC) of the Asan Institute for Life Sciences, Asan Medical Center. The IACUC abides by the Institute of Laboratory Animal Resources (ILAR) guide. All animals were given water and feed ad libitum and were housed under stable air and humidity conditions. Nine male Sprague–Dawley (SD) rats weighting 250–300 g (Ja Bio, Suwon, Republic of Korea) were used for the experiments, three of which died for unknown reasons during follow-up. We defined the time point at which the surgical procedure (LN dissection) and radiation were performed as the origin point (0W) (Figure S1A). Swelling and the lymphatic drainage patterns in NIRF-ICG lymphangiography confirmed whether lymphatic obstruction had occurred in the dissection limbs at one week after 0W (Figure S1B). Only valid animals were used for follow-up analyses, all others were sacrificed. The first measurement was started after verification of model formation (1W). All measurements were performed at daily intervals until 2^nd^ weeks (2W), and at weekly intervals until week 7 (7W) after origin point. The follow-up data obtained at each measurement comprised ICG fluorescence images, signals and limb volumes. Static and dynamic analysis were performed on the data obtained through imaging and signal processing (Figure S1A). During the follow-up analyses, the animals were housed in separate cages. At the end of the experiments, all animals were euthanized using carbon dioxide gas for 7 minutes, in accordance with the American Veterinary Medical Association guidelines.

### Surgical and radiation procedures for the lymphedema animal model

The model rats were not fed during the day before surgery, although water was still provided. The animal procedures were performed on specific premises with specific instrumentation such as a surgical microscope system (F170, Carl Zeiss, Germany), various microsurgical instruments and an electrocautery (Symmetry Surgical Inc., TN). All animals were operated on by the same researcher. The animals were preoperatively anesthetized with tiletamine/zolazepam (50 mg/kg; Zoletil, Virbac, France) mixed with xylazine (volume ratio 5:1, Rumpun; Bayer Korea, Seoul, Republic of Korea) after being induced with 4% isoflurane gas. After anesthetization, the fur in the upper limb was shaved with electric clippers and depilatory cream. Subsequently, 0.05-mL Evans blue solution (30 mg/mL solution in 0.9% saline; Sigma, St Louis, MO) was subcutaneously injected into the left palm to visualize the lymph vessels and LNs. The skin of upper limb and proximal skin were then separated by blunt dissection by following an incision line along the circumference of the upper limb based on the neck boundary and axillary area. After this skin blunt dissection, the blue-stained (because of Evans blue) axillary lymph nodes (ALNs) could be identified between the lateral border of the pectoralis major and the dorsal side of the latissimus dorsi in the ventral position. The ALNs and surrounding adipose tissue were carefully removed by sharp dissection. The brachial lymph nodes (BLNs) with the blue dye and adipose tissue surrounding them are located in the space between the triceps brachii and latissimus dorsi. Similar to ALNs, the BLNs were removed carefully by sharp dissection (Figure S2A). After the LN dissection, the cut edge of the skin was cauterized by a portable electrocautery unit (Bovie^®^ #430102; Symmetry Surgical Inc., Antioch, TN) and both ends of the skin were fixed to the muscles with an interval of approximately 1 mm (gap suture) to maintain disconnection of the intradermal lymphatic vessels (Figure S2B). Gentamicin (0.08mg/kg; Gentamicin Inj.; Shin Poong Pharm., Seoul, Republic of Korea) was injected to prevent bacterial infection after the surgery and ketoprofen (1 mg/kg; SCD Ketoprofen Inj.; SamChunDang Pharm., Seoul, Republic of Korea) was injected intramuscularly each week at the surgical area to reduce pain and inflammation.

The surgical area in the dissection limb was irradiated to provoke tissue fibrosis at one day postoperatively. The animals were anesthetized prior to being irradiated with a 20-Gy single dose using an X-Rad 320 device (Precision X-Ray, Madison, CT). The radiation treatment was only applied in the surgical area with the rest of the body protected with an 8-mm lead plate covering (99% radiation shielding). The total radiation dose was delivered in 10 fractions at a rate of 1 Gy/min to reduce the risk of morbidity.

### NIRF imaging system and ICG solution

An NIRF imaging system was utilized for measuring accurate lymphatic flow in the experimental rats. This customized system was modified with a commercial digital camera (Cannon EOS 50D; Cannon, Tokyo, Japan) for NIRF-ICG lymphangiography. A modified CMOS sensor was used to obtain the NIRF-ICG signal by removing the low pass filter in front of the sensor and installing a bandpass filter (FF01-832/27-50-D; Semrock, West Henrietta, NY) in front of the macro lens, which isolates the NIR spectrum from 810 nm to 850 nm. The bandpass filter was selected considering the emission peak of the fluorescence spectrum (828 nm) of the injected ICG solution. The ICG dye was excited by a light source with a total output power of 3.72 watts consisting of 12 high-power LEDs with a 730-nm peak (LST1-01G01-FRD1-00; Opulent Americas, Raleigh, NC) (Figure S3). The frequency spectrum was measured by a commercial visible-IR spectrometer (USB-650 Red Tide Spectrometer; Ocean Insight, Orlando, FL). The system was then able to obtain high-accuracy images with high spatial resolution. We tested the spatial resolution and the signal-to-noise ratio (SNR) of this modality using a lymphatic channel sheet (LCS) which is a bio-implantable artificial lymph vessel containing 250-μm microfluidic channels [^22^]. The signal-to-noise ratio was 14.8 decibels for background and 10.7 decibels for near-channel area in the resulting NIRF-ICG images obtained with this process (Figure S4A). In spatial resolution analysis, the diameter of the channels in the images (271 μm, 254 μm), which was calculated from full width at half maximum (FWHM values) of the peaks, was almost identical to the actual diameter (250 μm) (Figure S4B). The system also obtained fluorescence images with a resolution in the 10 micron range under the epidermis (thickness of approximately 350∼450 μm, [^27^]) Because the diameter of collecting lymphatics is 100∼150 μm in the rat [^28^], our system proved suitable for analyzing the micro-lymphatic system in small animals.

### NIRF-ICG lymphangiography and imaging processing

NIRF-ICG imaging was used to analyze the static (lymphatic drainage pattern and lymphatic leakage) and dynamic (lymphatic contraction or lymphatic pump) changes in the disrupted lymphatic flow. Briefly, ICG medicine powder was dissolved in a 25 mg/ml bovine serum albumin (BSA) solution at a concentration of 20 μg/ml because ICG molecules are stabilized by albumin and this complex improves the intensity of the NIRF-ICG signal [^29,30^]. After the animals were anesthetized using tiletamine/zolazepam and xylazine mixture (volume ratio 10:1, not same ratio with surgical procedures), the fur on the upper limbs was shaved and treated with a depilatory cream prior to each measurement to prevent scattering of the NIR light. We did not use isoflurane gas anesthesia in the NIRF-ICG imaging protocol because it can affect lymphatic vessel contractility [^31^] An ICG-BSA complex solution of 6 μL was injected slowly into the intradermal layer between the toes of the palm using 34-gauge needles. Real-time images were acquired after confirming that the ICG dye had stably spread throughout the entire lymphatic vessel network in the upper limbs (usually 3 minutes after injection) (Figure S3). In the static analysis, NIRF-ICG image data were obtained from the dorsal position of upper limbs where the collective lymph vessels were identified.

In the dynamic analysis, the ICG intensity was measured in the distal collective lymph vessels at a point 2 mm away from the wrist, the region of interest (ROI), using NIRF-ICG video images (Movie S1, Move S2). To minimize the effects of injection pressure, the measurement was not started until 3 minutes after the first ICG solution injection. The camera system originally offered 30 fps, but we converted the sampling rate to 3 Hz due to the image denoising process of smoothing the signals. Because changes in the NIRF-ICG signal intensity arise from robust contractile movement of lymphangion (the functional unit of a lymph vessel), in vivo signals from lymphatic contraction can be obtained in NIRF-ICG lymphangiography. After converting them to time-domain signals, the cubic spline interpolation which is usually used with electrocardiogram (ECG) signals was applied to remove background noise from the in vivo signals of lymphatic contraction [^32,33^]. Because the fluorescence intensity does not relate to the ICG molecular conditions inside the lymph vessels and the difference between systolic and diastolic points is important for lymphatic contraction [^19^], the signals were normalized after the baseline correction (Figure S5). In order to identify frequency components of lymphatic contraction, we used the FFT algorithm expressed by Formula 1:

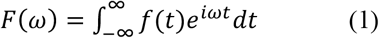

where *F*(*ω*) is the transformed signal in the frequency domain; *f*(*t*) is the original time-domain signal; *i* is the imaginary number; *ω* is the circular frequency variable; and *t* is the time variable.

We next applied the CWT algorithm to investigate the characteristics of the lymphatic contraction in the time-frequency spectrum [^34,35^]. We utilized the well-known Morlet wavelet, which is a sine wave in Gaussian window, for this purpose and the CWT was represented by Formula 2 as follows:

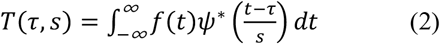

where *T*(*τ, s*) is the correlation coefficient compared with the wavelet; *f*(*t*) is the original signal; *ψ* is the wavelet (*denotes the complex conjugate); *τ* is the shift parameter of the wavelet; *s* is the scale parameter of the wavelet; and *t* is the time variable. The scale parameter *s* could be changed to the pseudo-frequency *F*_p_ in accordance with Formula 3.

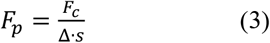

where *F*_p_ is the pseudo-frequency; *F*_c_ is the center frequency of the wavelet; and Δ is the sampling period. Images were processed using ImageJ 1.48v software (NIH, Bethesda, MD) and Matlab 2013a (MathWorks, Natick, MA). The FFT and CWT signal processing was performed using Matlab 2013 and the Origin Pro 9 (Origin 9.0; OriginLab, Northampton, MA) software for numerical computation and visualization.

### Limb volume measurement

The purpose of the volume measurements was to validate our lymphedema rat model during the follow-up period. Although swelling is not a direct indicator of lymph flow, it reflects the environment around this flow by indicating the retention of lymph fluid and tissue deformation. Volume measurements of the upper limbs on both sides were performed using photographic analysis. The animals were placed in the same dorsal position after anesthesia as shown in Figure S6. Similar to the protocols used in clinical practice, we assumed that the shape of the upper limb from the wrist to the elbow was a conical frustrum and calculated the upper limb volume using the frustrum approximation. The values measured using ImageJ software were the diameter of the wrist, the diameter from the cubitus, and the distance between the wrist and elbow point. The volume was then calculated using formula 4 below, which represents the conical frustum approximation:

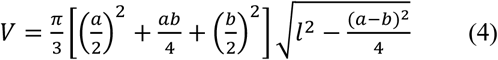

where *V* is the volume of each limb; *a* is the diameter of the wrist; *b* is the diameter of the elbow joint; *l* is the superficial length from wrist to elbow; *h* is the height of the upper limb frustum.

### Statistical analysis

Statistical analyses were conducted using GraphPad Prism 9 (GraphPad Software Inc.; Boston, MA) and Microsoft Excel 2019 (version 2111; Microsoft Corporation, Redmond, WA). Differences between the groups were compared using the Student’s t-test and Mann-Whitney U test, and P values < 0.05 was considered to indicate a statistically significant difference. The data were expressed as mean values ± standard deviation. In the box-and-whisker diagram, the IQR represents the statistical dispersion in the boxplot, which is the spread range of the data. The IQR values were obtained by subtracting Q1 (the lower quartile, 25^th^ percentile) from Q3 (the upper quartile, 75^th^ percentile).

## RESULTS

### Volume measurements

There were no significant volume differences between the control and dissection limbs before the surgery and radiation (i.e. at the origin point, 0W). However, the volume in the dissection limbs were found to be significantly increased at 1W due to the onset of the acute lymphedema caused by the surgery and radiation. Subsequently, these volumes decreased continuously until the 2W time point as the surgical injury was recovered from. From the 2W to the 7W time points, the chronic lymphedema condition was maintained, and the volume increased by about 12% or more in comparison to the control limb. The volume differences between the control and dissection limbs were all statistically significant other than at 11D (*P* < 0.05 and *P* < 0.01). As shown in Figure 1, the median values of volume in the dissection limbs were 1626.8 mm^3^ compared to 1435.0 mm^3^ in the control limbs, which was a significant difference throughout the follow-up period (*P* < 0.001) indicating that swelling had been maintained in the dissection limb.

**Figure 1.**
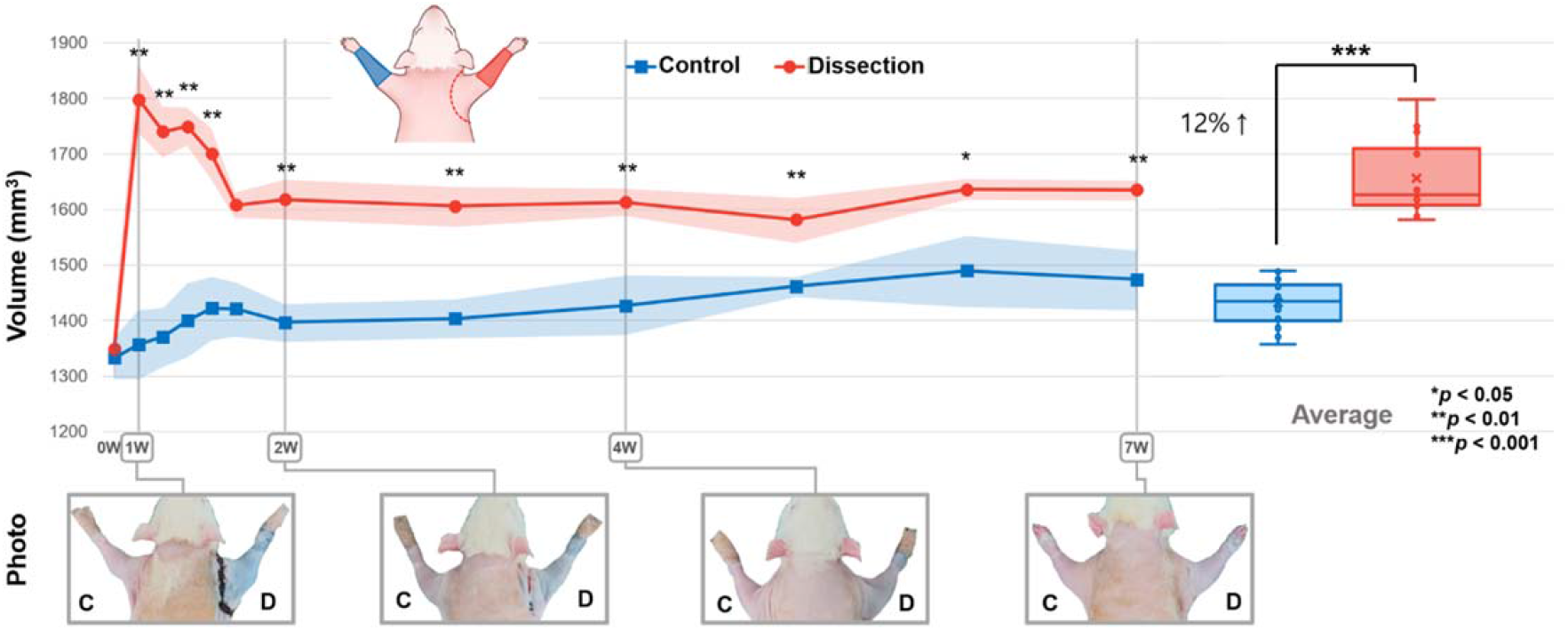
The volume changes in both the control (“C”) and dissection (“D”) rat limbs during the follow-up period. The condition of the animals just prior to surgery and radiation was denoted as “0W”. The volume of the dissection limb increased by approximately 12% compared to the control group, indicating a significant difference between the two groups. Through the whole follow-up period, the volume of the dissection limb (median value: 1632 mm^3^) was significantly different from the control group (median value: 1424 mm^3^) as determined by the Mann-Whitney U test (\**P*<0.05, \*\**P*<0.01, and \*\*\**P*<0.001). The LN dissection and radiation had a significant effect on the limb volume of the animals and this effect persisted throughout the follow-up period.

### Static analysis of lymphatic drainage patterns

Lymphatic drainage pattern analysis via NIRF-ICG lymphangiography is commonly used to determine the severity of lymphedema in clinical practice. Figure 2A shows the changes we here observed in the lymphatic drainage pattern of both upper limbs in the rats over time (1W, 2W, 4W, and 7W). The patterns in the controls showed normal lymph drainage in the upper limb. Lymph fluid in the distal region is collected to the collective lymphatic vessels and moves along the vessels to the BLNs (round white area). In the dissection limbs however, different patterns from those observed under normal conditions were evident throughout the upper limb areas in the model animals. To quantify the differences between the control and dissection limbs, the lymphatic drainage patterns in NIRF-ICG lymphangiography were classified as linear (normal), or as splash, stardust, or diffuse (abnormal, in order of low severity), based on established clinical standards [^36–38^]. The frequency of each pattern type in the proximal region was counted because that area is relatively larger than the distal region and they can be more clearly distinguished. Figure 2B shows images used in this study for pattern classification and the quantitative results at each follow-up time point. In the control group, only a normal lymph drainage pattern (linear) was observed over the entire follow-up period. In the dissection limbs however, stardust and diffusion patterns were predominant after surgery and radiation treatment due to the removal of BLNs and lymphatic flow disconnection. After 4 weeks (4W), more than half of the patterns were splash type. This indicated a relatively mild level of severity only but they never returned to a linear form.

**Figure 2.**
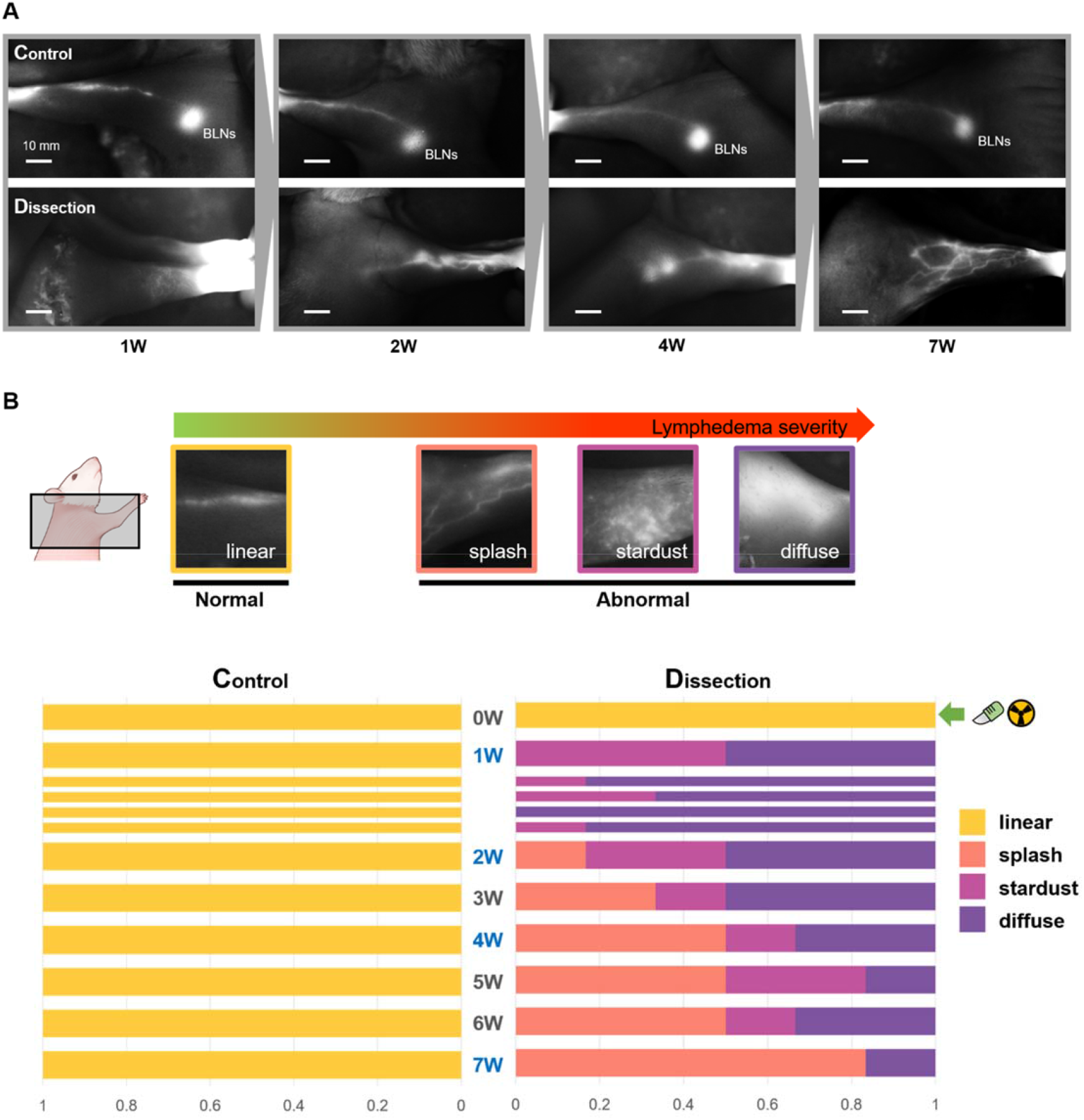
(**A**) The changes in lymphatic drainage pattern in NIRF-ICG lymphangiography. To ensure that the analysis was conducted in a consistent and unbiased manner, all of the image data used in this study were obtained from the animals being placed in identical positions for both the control and dissection groups. (**B**) The representative images for classification of patterns and their normalized quantified results in the follow-up period from 0W to 7W. Based on the clinical standard, the patterns were classified as one normal pattern (linear) or three abnormal patterns (splash, stardust, and diffuse). In cases where multiple patterns were observed, the pattern with the widest range was counted and quantified for analysis.

### Static analysis: leakage of lymph fluid

The disruption of the lymphatic flow increases the pressure in the collective lymph vessels which leads to lymph fluid leakage into the interstitial tissue through the endothelial-intercellular junction [^39,40^]. This is another cause of worsening lymphedema and is different from postsurgical lymphatic leakage It and can be identified through static analysis of NIRF-ICG lymphangiography images. We determined the degree of lymphatic leakage in our present analyses by measuring the ICG intensity in a section containing collective lymph vessels at a point 10 mm from the wrist in the control and dissection limbs. A cross-section of the ICG intensity revealed a peak-shaped spectrum form centered on the collective lymph vessels, with the thickness of the peak indicating how much the ICG dye had spread around the lymph vessels (Figure 3A). The graph in Figure 3A contains representative results from the control and dissection limbs. These data were quantified using the full width at half maximum (FWHM) of each peak and collected throughout the whole follow-up period. As shown in Figure 3B, the FWHM of the peak in the dissection limbs was increased by approximately 2.65-fold compared with the control limbs, which was a significant difference (*P* < 0.001). The median value was 2.14 mm in the control limbs and 5.69 mm in the dissection limbs.

**Figure 3.**
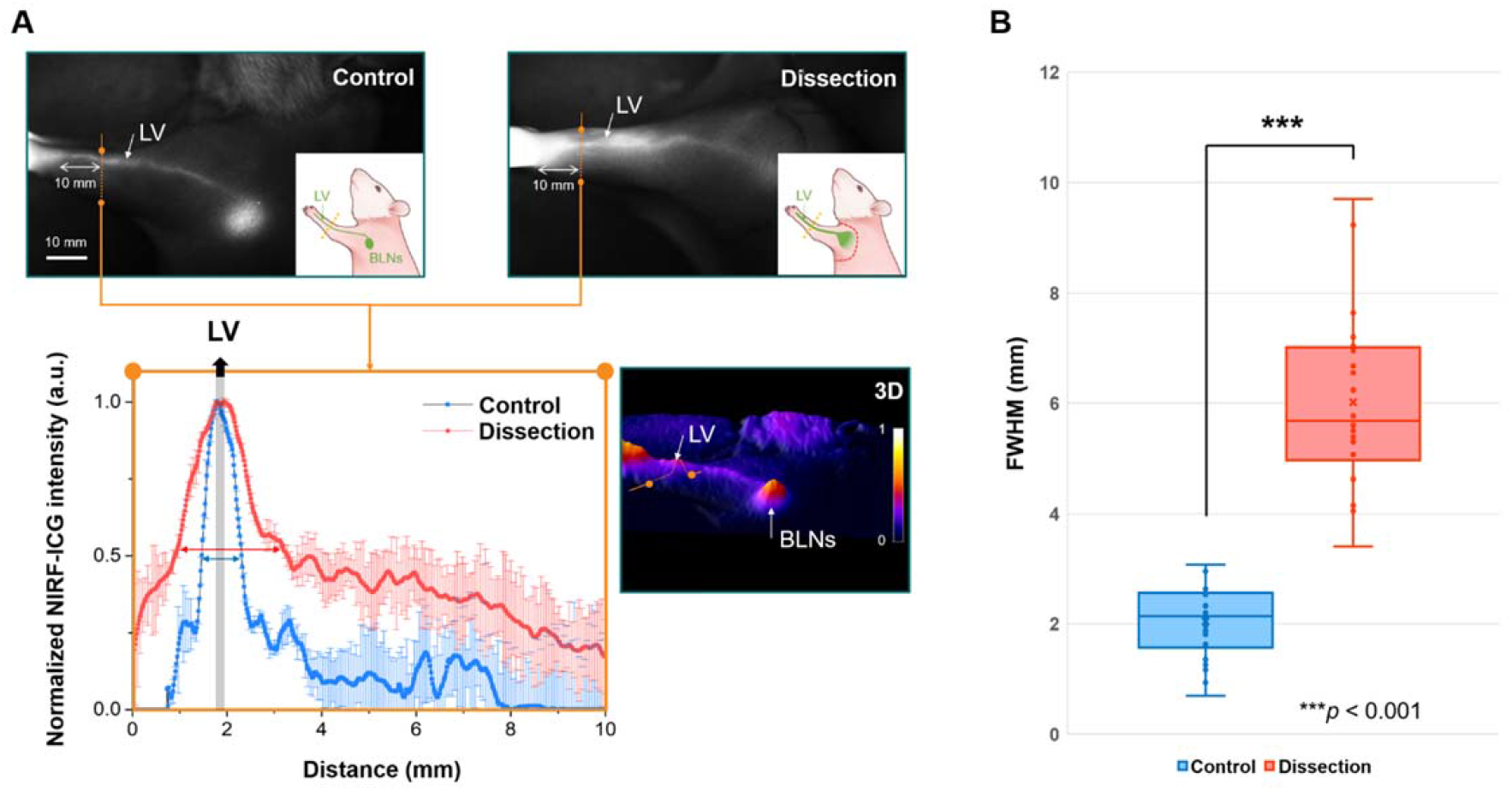
Leakage of lymph fluid around the distal collective lymphatic vessels (LV) in the dissection limb observed on NIRF-ICG lymphangiography. (**A**) The representative NIRF-ICG images and the corresponding graph of normalized ICG intensity in both the control and dissection limbs which was obtained from the images. The ICG intensity was measured at a point 10 mm from the wrist perpendicular to the vessels and the data was averaged three times. The peak in the graph is the LV location (gray area). The graph was corresponded with the cross-sectional ICG intensity data obtained from the 3D image at the measurement point of the NIRF-ICG images. The orange dots and the lines between them represent the same points on the image and graph. (**B**) The box plots representing the full width at half maximum of the ICG intensity peaks in the graph around the lymph vessels in the control and dissection limbs in the follow-up period. The FWHM values present the thickness of the ICG intensity peak. The results for the lymphatic leakage around the vessels were significantly different between the groups (\*\*\**P* < 0.001).

### Dynamic analysis: time-domain signals of lymphatic contraction

Figure 4A shows the representative NIRF-ICG imagery and ROIs used to measure in vivo lymphatic contraction in the control and dissection limbs. The rats were anesthetized by xylazine injection and hence had resting skeletal muscles [^41^] so that the factors that affect lymphatic contraction were minimized. Because the lymphatic muscle cells try to maintain a robust contraction [^42,43^], the signals of lymphatic contraction, which were measured by NIRF-ICG lymphangiography, usually presents as periodic waveforms in the time-domain. We obtained pulse-shaped waveforms from the ROIs of distal lymph vessels in each control and dissection limb (Figure 4B). According to previous research results observed in vivo and in vitro [^44^], a lymphangion repeats three phases which are the inflow of lymph fluid (phase 1), expansion of the lymphatic vessel wall (phase 2), and the drainage of lymph fluid (phase 3). This three-phased movement was observed in the pulse shape of the lymphatic contractile waveform in the control group. By contrast, the contractile pulse shape was altered in the dissection limbs because of lymphatic disruption in the proximal area. Because the movement of lymph fluid by the contraction was not efficient, the peak shape of the pulse (from phase 2 to phase 3) becomes blunt and the distinction between systole and diastole becomes unclear. The lymphatic disruption also changed the frequency of lymphatic contraction. Figure 4C presents the number of lymphatic contractions per 5 minutes during the whole follow-up period. The average frequency of lymphatic contraction (i.e. the average number of contractions completed every 5 minutes) was 6.28/min in the control limb was increased to 9.92/min (about 58% increase) in the dissection limb. The scatter of data increased in the dissection limb compared to the control limb. The IQR in the control limb group was 5.5 while that in the dissection limb group was 14.75. The number of lymphatic contractions in both groups were significantly different (*P* < 0.001), withs the average values measured at 31.4 in the control limbs and 49.6 in the dissection limbs. The median value was 32 in the control limbs and 48.5 in the dissection limbs.

**Figure 4.**
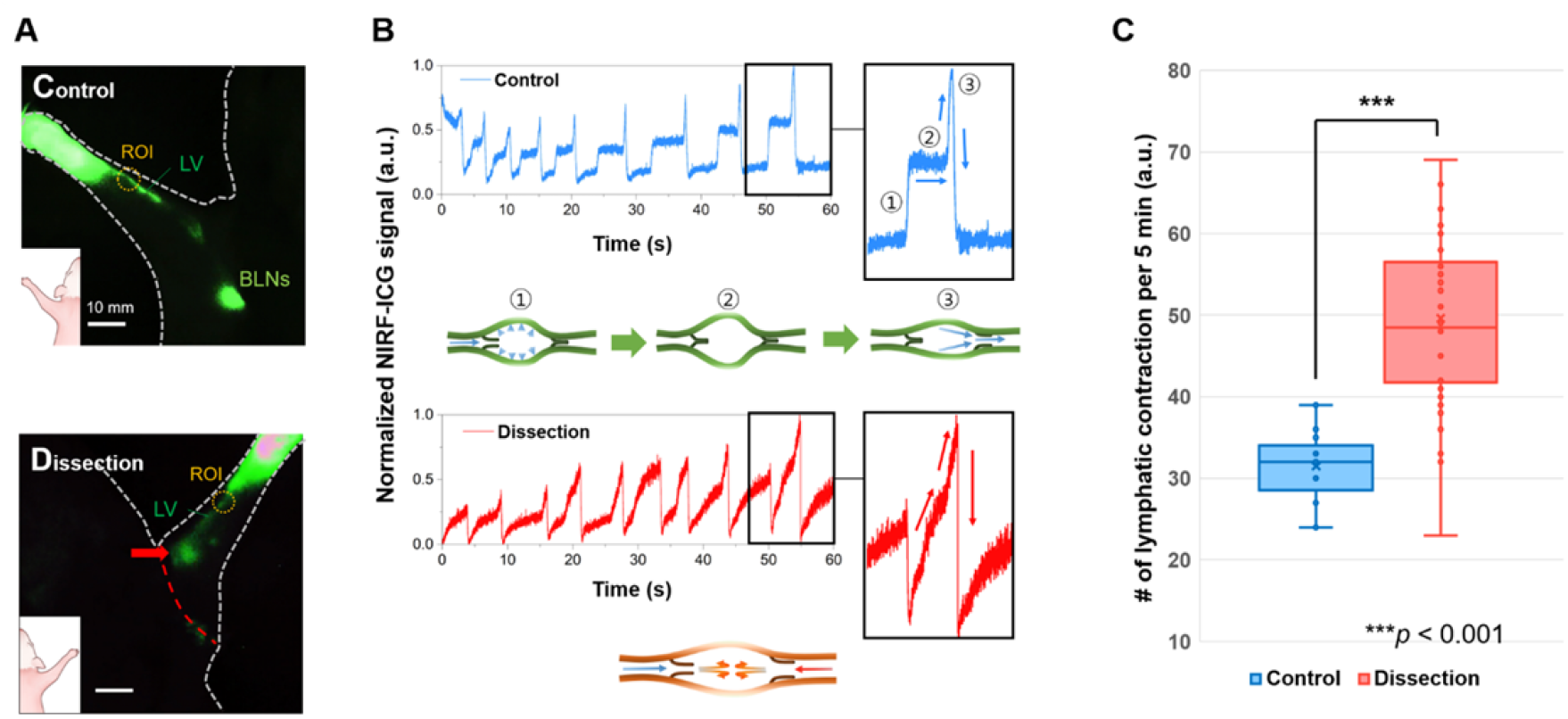
(**A**) The ICG images and region of interest (ROI) used to obtain the signals for lymphatic contraction in the control and dissection limbs. The white and red dash lines represent the body shape and the incision line in the dissection limb, respectively. Additionally, the red arrow indicates the diffusion of lymph fluid in the proximal area near to the incision line. (**B**) The representative lymphatic contractile waveforms in the distal lymph vessels of both groups for 60 seconds (1 minute). The y-axis indicates the normalized NIRF-ICG signals for each ROI. In the control limbs, the lymphangion maintains contraction by repeating the phases of ①lymph fluid inflow, ②lymphatic vessel wall expansion, and ③lymph fluid drainage, and the pulse shape represents these three phases. In the dissection limb, the contractile pulse shape was changed compared to the control limbs because of lymphatic disruption in the proximal area. The overall peak shape of the pulse became blunt, and there was a loss of distinction between the systolic and diastolic phases. (**C**) The box plots for the number of lymphatic contractions per 5 minutes in each group during the follow-up period. The average value (31.4 vs. 49.6) and interquartile range (IQR) of the lymphatic contractions were increased significantly in the dissection limbs (\*\*\**P*< 0.001).

### Dynamic analysis: frequency spectrum of lymphatic contraction

We converted the time-domain wave form data collected for 5 minutes from the ROIs to a frequency-domain spectrum to investigate the frequency characteristics of the signal using FFT signal processing with a 5 mHz frequency resolution. If there had been a regularity of lymphatic contraction, its frequency would be represented by the peaks in the frequency-domain spectrum (Figure 5A). In the frequency domain, the most meaningful spectrum was found to exist below 1 Hz. This indicated that in vivo lymphatic contraction usually moved at less than a 1-Hz period. As shown in Figure 5B, we observed a specific peak between 0.08 Hz and 0.13 Hz in the frequency domain spectrum of the control limbs during the whole follow-up period. However, these frequency peaks were not present in the dissection limbs. To compare these FFT results, the same lymphatic contraction data were analyzed using CWT signal processing. The magnitude of the correlation coefficient with the wavelet, which is represented by a color bar (red: higher correlation, blue: lower correlation), indicates the specific frequency in the lymphatic contraction signal over time. Figure 6 shows representative CWT results from the control and dissection groups. We found a red-color frequency band at around 0.1 Hz over time in the control limb results, which was located in a frequency spectrum that was similar to the frequency peak of the FFT (Inset, Figure 6). The frequency peak of the FFT and the frequency band of the CWT represented the periodic movements of lymphatic contraction. In the dissection limbs, a specific frequency band could not be identified as a result of the disappearance of the frequency peak in the FFT. This indicated that there was an irregular contractile movement of distal lymph vessels in the dissection limb.

**Figure 5.**
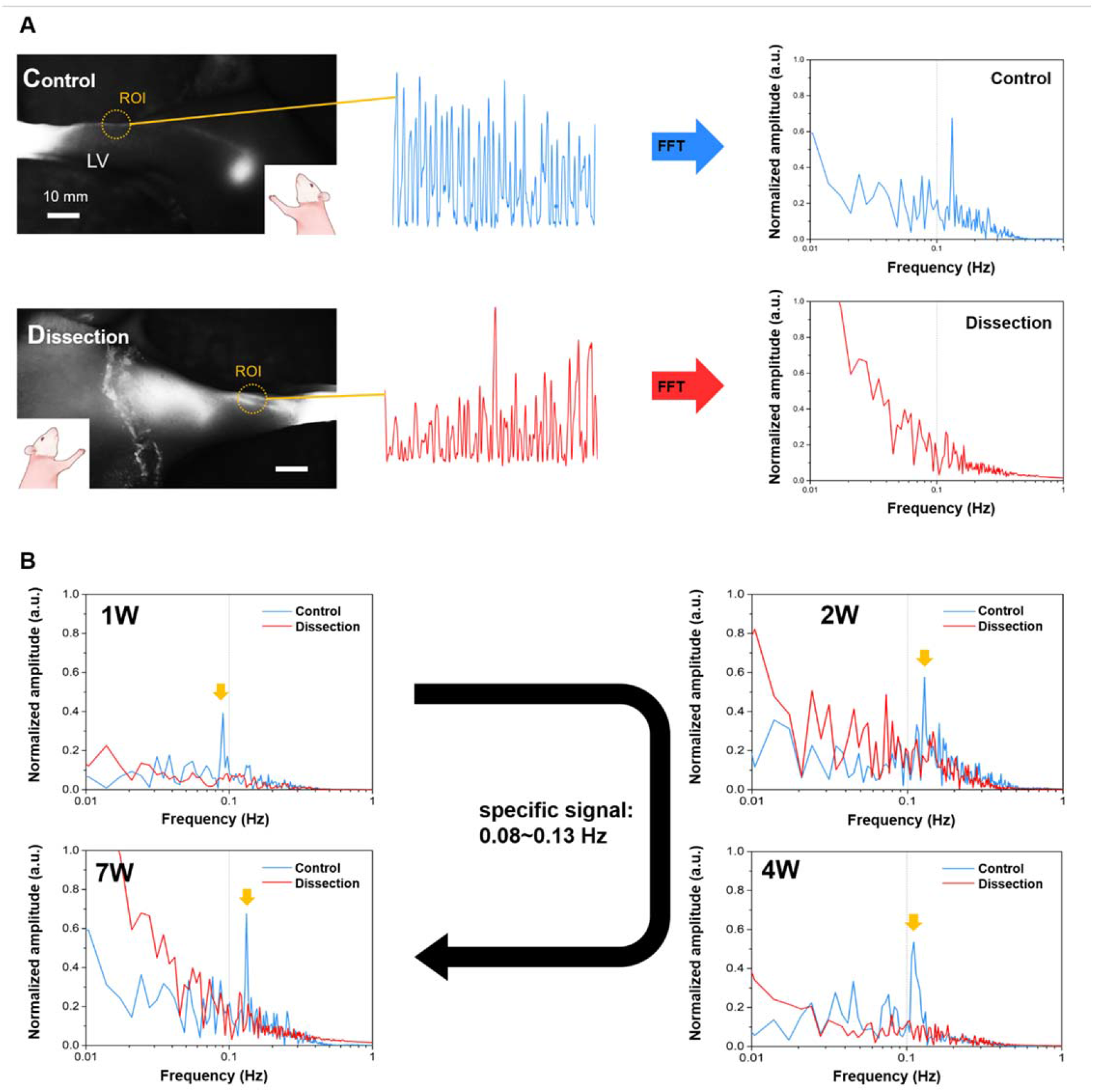
The FFT results of each lymphatic contraction in the control and dissection limbs. (**A**) The conversion of time-domain waveforms of lymphatic contraction in the distal lymph vessel (ROI), obtained from real-time NIRF-ICG video, into frequency-domain spectra using FFT signal processing. By performing the signal processing, the periodic component of the signal became visible as distinct peaks in the graph of the frequency spectrum. (**B**) The comparison of FFT signals between the control and dissection limbs at 1W, 2W, 4W, and 7W of the follow-up. The frequency-domain spectra were plotted with the logarithmic scale of the frequency on the horizontal axis and the normalized amplitude of the signal on the y-axis. Upon analysis, a specific frequency peak was observed between 0.08 Hz and 0.13 Hz in the control group only (indicated by the yellow arrow).

**Figure 6.**
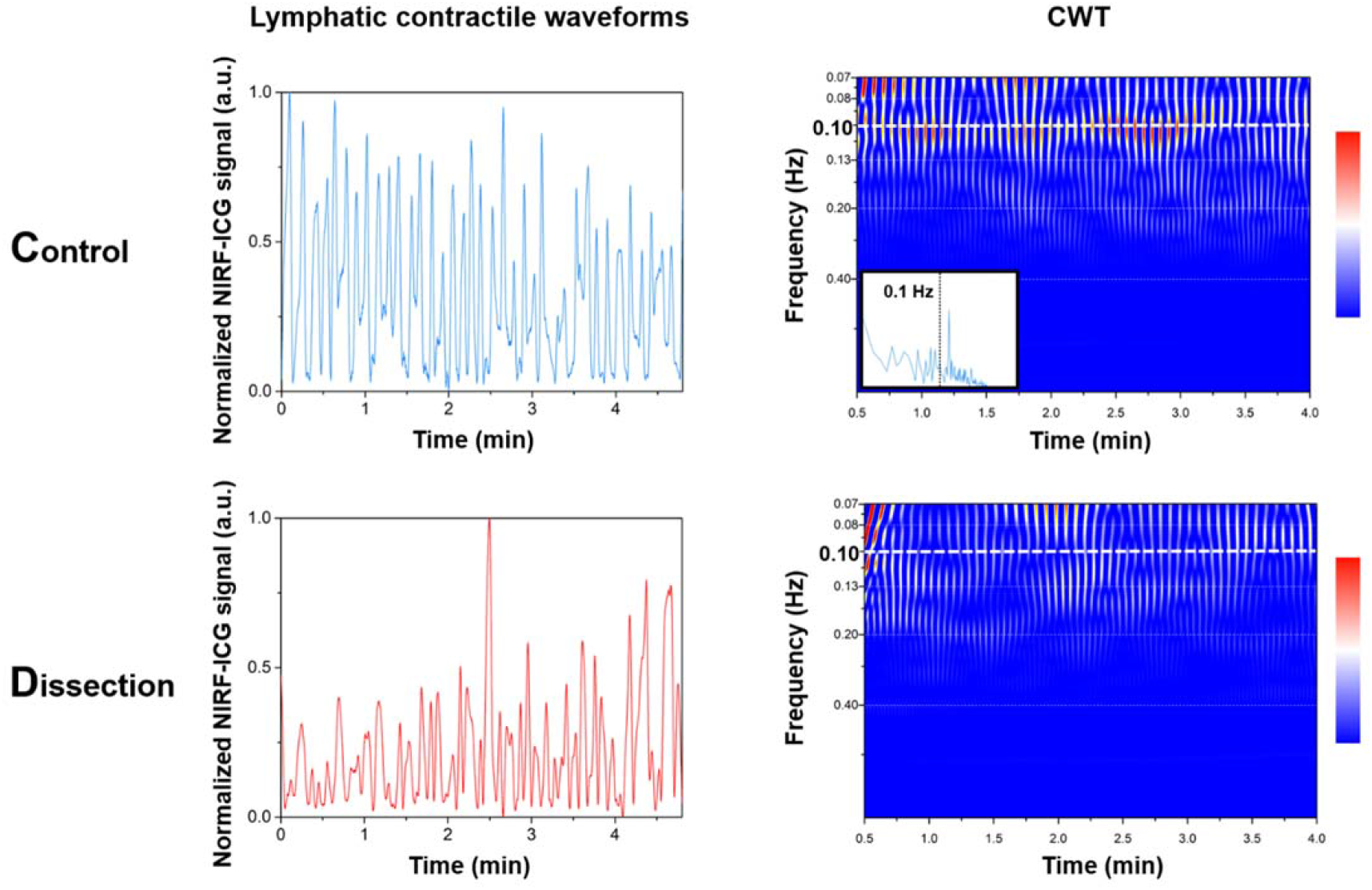
The results of the Continuous Wavelet Transform (CWT) analysis based on the lymphatic contractile waveforms with the magnitude of a specific frequency in the lymphatic contraction signal at a particular time represented by a color bar. The red color indicates a greater magnitude of the specific frequency. In the control group, the frequency band was observed at around 0.1 Hz, which was found to be consistent with the FFT result (the inset). The results of CWT signal processing provides additional evidence for the presence of a consistent frequency component in the lymphatic contraction waveform at 0.1 Hz in the control limbs and it was faded away in the dissection limbs.

## DISCUSSION

The lymphatic system maintains a robust homeostasis of lymphatic contraction via chemical feedback from the Ca2+ and nitric oxide (NO) levels within the lymphatic muscle cells (LMCs) when there are no external factors affecting these conditions [^28,45,46^]. Hence, robust lymphatic contraction under regular conditions has been reported in several studies including that of Dixon et al. [^47–50^]. However, under lymphedema conditions, the homeostasis of lymphatic contractions is disrupted. Liao et al. demonstrated that lymphatic contraction is attenuated by molecular regulation abnormalities under the inflammatory conditions of lymphedema, and provided important insights into the dysfunction of lymphatic contraction in this disorder [^51^]. In our present study of a lymphedema rat model, an irregularity of lymphatic contraction caused by lymphatic disruption in the proximal area was observed at the collecting lymph vessels in the distal area. This finding indicated that a disruption to lymphatic contraction may occur due to physical or mechanical causes as well as inflammatory conditions. According to a recent study by Choi et al., sustained lymphatic flow involves not only the formation of lymphatics but also the maintenance of lymphatic valves by the Piezo1 factor, which is a mechanosensitive ion channel protein [^52^]. Because the function of lymphatic valves plays an important role in maintaining the regularity of lymphatic flow, the disturbance of this flow leads to lymphatic dysfunction. Because the frequency of lymphatic contraction can be altered by small pressure fluctuations [^43^], a disruption to the lymphatic circulation may have a far wider range of effects than expected. Indeed, we here observed that a disturbance to the lymph flow in the proximal area caused changes in the lymphatic function in the distal area, and also that the regularity of lymphatic contraction was broken. Moreover, this irregularity of lymphatic contraction was detected through the dynamic analysis of NIRF-ICG lymphangiography (NIRF-ICG lymphangio-dynamics) in-vivo, and presented as the absence of specific frequency signals.

Based on its clinical utilization, the use of in vivo NIRF-ICG lymphangiography to date has principally focused on the static analysis component (mainly lymphatic drainage pattern [^53^]). This method provides spatial information and can be used to assess the current condition of the lymph vessels such as the distributed lymphatic drainage pattern or the expansion of the lymphatic vessels. This information makes it possible to determine the leakage of lymph fluid from the lymphatic vessels to the surrounding tissue, and enables a diagnosis of lymph circulation disorders to be made based on the abnormal distribution of lymph fluid. In addition, this modality enables the location of the lymph vessels to be identified for the purpose of functional microsurgery such as lymphovenous anastomosis surgery (LVA), which constructs a lymphatic bypass to a nearby vein. On the other hand, the state of lymphatic contraction (i.e. its degree of regularity) is important for dynamic analysis. Because the lymphatic contraction signals are produced by the rhymical movement of LMCs and by the valve function in the lymphangion, problems in lymphatic system can be predicted through the use of dynamic analysis component of NIRF-ICG lymphangiography. The NIRF-ICG lymphangiography for assessing lymphatic function and circulation is similar to the electrocardiography (ECG) for identifying problems in the heart and cardiovascular system. The lymphangion, which provides a driving force for lymphatic circulation, plays the same role in lymphatic circulation with the heart in the cardiovascular circulation. In both techniques, abnormalities in the signals can indicate problems with circulation system, although the difference is that ECG uses electrical signals in the heart while NIRF-ICG lymphangiography uses optical signals. The similarities between the two techniques highlight the importance of non-invasive diagnostic modalities in identifying and diagnosing abnormalities in the cardiovascular and lymphatic systems.

Figure 7 shows a comparison between static (quantified lymphatic drainage pattern and leakage of lymph fluid) and dynamic analysis (lymphatic contraction in FFT and CWT) at the 1W, 2W, 4W, and 7W follow-up time points. The volume data indicates the difference between control and dissection limbs at the same periods. Prior to LE formation (0W), there was no significant difference between the control and dissection limbs of the model animals, and the static analysis of NIRF-ICG lymphangiography revealed a linear pattern of collecting vessels because the lymph fluid was concentrated in these vessels via normal lymphatic drainage. In the dynamic analysis, FFT results showed a specific frequency peak around 0.1 Hz, and CWT results revealed a specific band at the same frequency due to the regularity of lymphatic contraction under stable anesthesia. After the LN dissection and radiation however, the static and dynamic analysis results changed dramatically with the volume differences. The damage and disruption caused to the lymphatic flow in the proximal area induced lymphatic leakage from the collecting lymphatic vessels in the entirety of the upper limbs, leading to acute swelling and an abnormal patten of lymphatic drainage. This abnormal lymphatic environment affected the dynamic movement of the lymphatic contraction and its regularity, even in the distal area, and the characteristic frequencies that could be observed under normal conditions were no longer present in the FFT and CWT results. In our volume evaluations and static analysis, disruption to the lymphatic circulation could be identified by the relative values obtained in the lymphedema conditions compared to the valve functions under normal conditions. However, the dynamic analysis of NIRF-ICG lymphangiography allowed us to identify the lymphatic circulation disruptions via the presence or absence of a specific frequency signal, and could provide therefore more objective evaluation criteria.

**Figure 7.**
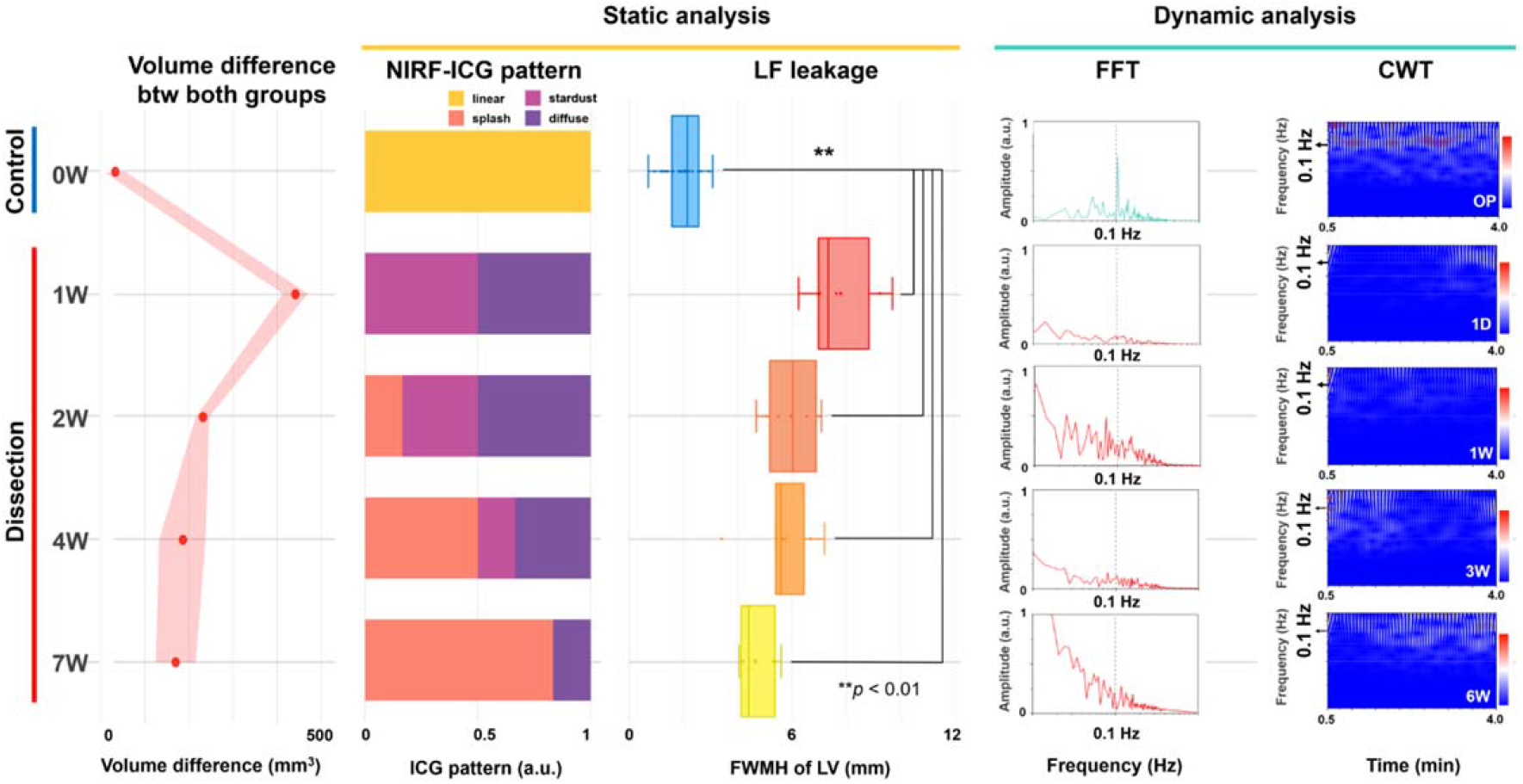
The volume differences between the control and dissection limbs and the results of ICG analysis using static (ICG pattern and LF leakage) and dynamic (FFT and CWT) methods (\*\**P* < 0.01) on the same time line of the follow-up period. The quantified results of the static analysis could be evaluated by comparing it to the relative value of the control group. In contrast to static analysis, the dynamic analysis of lymphatic contractions focuses on the characteristics of the frequency regularity in the lymphatic system, and evaluates the presence or absence of regular contractions at certain frequencies (around 0.1 Hz in this study) to identify the disruption of lymphatic circulation.

There are several advantages to the use of dynamic analysis of NIRF-ICG lymphangiography in clinical practice. First, this modality shows potential as an early diagnostic method for lymphedema though the detection of disrupted lymphatic function before the damage spreads more widely. Our present findings indicate that proximal lymphatic damage affects the function of intact lymphatic vessels in the distal area. The evaluation of lymphatic function in this way after LN dissection or radiotherapy, both of which can damage the lymphatic system, may therefor predict the development of lymphedema. Second, because this dynamic analysis can be used to select lymphatic vessels with good function, the prognosis from functional surgery for the treatment of lymphedema such as LVA may be improved by using lymphatic vessels that remain well functioning [^40^]. If damaged lymphatic vessels are used for LVA, blood flow back into the lymphatic vessel may occur due to a lower pressure in these vessels and outflow obstruction [^54^]. Finally, the static analysis methods require an imaging system whereas the dynamic analysis methods only need optical detectors to collect NIRF-ICG signals over time. This will enable faster and more convenient analysis in a real-world clinical setting.

There were some limitations of this study of note. We did not confirm whether dynamic analysis could observe the development of lymphedema earlier than static analysis because the animal model was an acute rather than a progressive lymphedema model. Next, we did not provide information on how far from the site of LN dissection or radiation the lymphedema condition could be observed through the dynamic analysis due to this being a small animal study. Finally, the ROI in the distal area was a simple one-way structure directly connected to the site at which LN dissection and radiation were performed, but the human lymphatic network is far more complex than that of small animals.

In conclusion, the characteristic signals from lymphatic contractions under normal lymphatic conditions show a regular frequency of approximately 0.1 THz. We demonstrated that the signal change, which is especially interruption of the regularity of lymphatic contraction, may be an indicator of lymphatic disruption through animal model experiments through comparison with the results of static analysis. Although the static analysis of NIRF-ICG lymphangiography is a valuable diagnostic method, its dynamic analysis provides additional information that cannot be obtained from the static analysis alone. By performing the dynamic analysis of lymphatic circulation, such as assessing the pattern or regularity of the lymphatic contraction, it may allow detecting of abnormalities of lymphatic circulation in the early stages, enabling earlier intervention and treatment. Furthermore, incorporating the dynamic analysis into routine clinical practice may enhance the effectiveness of NIRF-ICG lymphangiography by improving its diagnostic accuracy and providing more detailed information about lymphatic circulation.

## ACKNOWLEDGMENTS

This work was supported by a grant (2021IL0035) from the Asan Institute for Life Sciences, Asan Medical Center, Seoul, Korea, and from the National Research Foundation of Korea (NRF) grant funded by the Korea government (Ministry of Science and ICT, MSIT) (No. NRF-2019R1A2C1009055, No. NRF-2021R1F1A1056527). We thank the core facilities of the Comparative Pathology Laboratory and Animal Experiment Laboratory at the ConveRgence mEDIcine research center (CREDIT), Asan Medical Center, for the use of their shared equipment, services, and expertise. The illustrations used in the figures were created by H. Cheon with the help of the medical contents Center of Asan Medical Center. We acknowledge Dr. Maria Nessa Gelvosa for assistance with the animal procedures.

## COMPLIANCE WITH ETHICAL STANDARDS

All animal procedures in this study were reviewed and approved by the Institutional Animal Care and Use Committee (IACUC) of the Asan Institute for Life Sciences, Asan Medical Center, in compliance with the Institute of Laboratory Animal Resources (ILAR) guide.

